## Supplementary Information for "Bridging polarities in metabolomics: Cross-ionization mode chemical similarity prediction between tandem mass spectra"

[1] Bioinformatics Group, Wageningen University & Research, 6708 PB, Wageningen, the Netherlands
[2] The National Phenome Centre, Section of Bioanalytical Chemistry, Division of Systems Medicine, Department of Metabolism, Digestion and Reproduction, Faculty of Medicine, Imperial College London, Hammersmith Hospital Campus, London, W12 0NN, United Kingdom
[3] Institute of Organic Chemistry and Biochemistry of the Czech Academy of Sciences, Prague, Czechia
[4] mzio GmbH, Bremen, Germany
[5] Centre for Digitalisation and Digitality (ZDD), University of Applied Sciences Düsseldorf, Düsseldorf, Germany
[6] Department of Biochemistry, University of Johannesburg, Auckland Park, Johannesburg, 2006, South Africa
* These authors jointly supervised this work.

### Supplementary Section 1. Hyperparameter optimization

#### a) All validation spectra

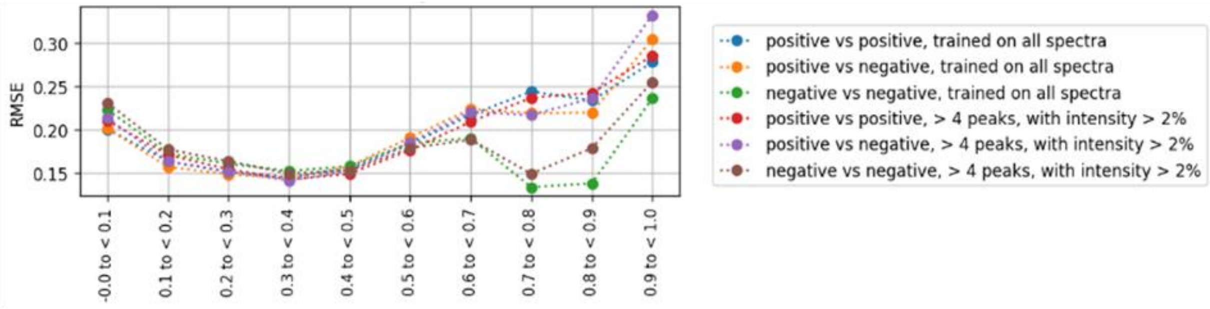


#### b) Validation set with > 4 peaks (intensity > 2%)

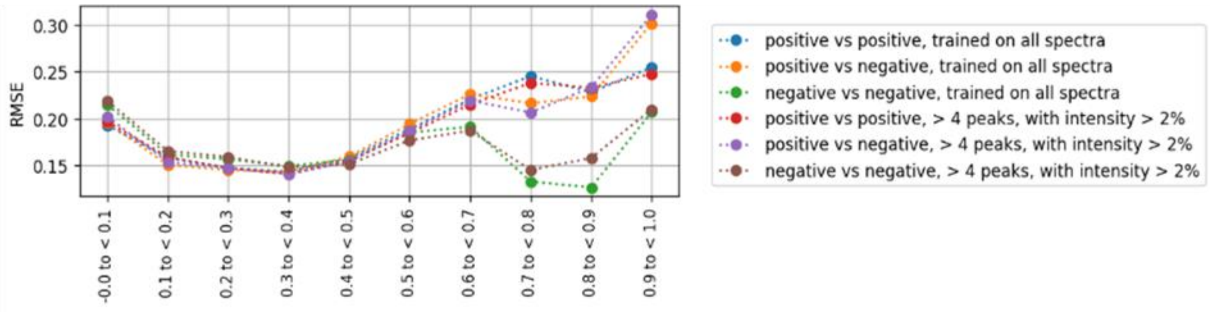


***Supplementary Figure 1.1: Average RMSE per bin for model trained with different filtering of training spectra.*** *For one of the models the training spectra only contained spectra that have at least 5 peaks at intensity >2%. The other model was trained on all spectra. The model trained on all spectra is used in the rest of the paper, since it resulted in best overall performance. Both models were trained with identical settings.* ***a)*** *Both models were validated with the standard validation set, containing all spectra.* ***b)*** *Both models were validated with the validation set only containing spectra with at least 5 peaks at intensity >2%.*

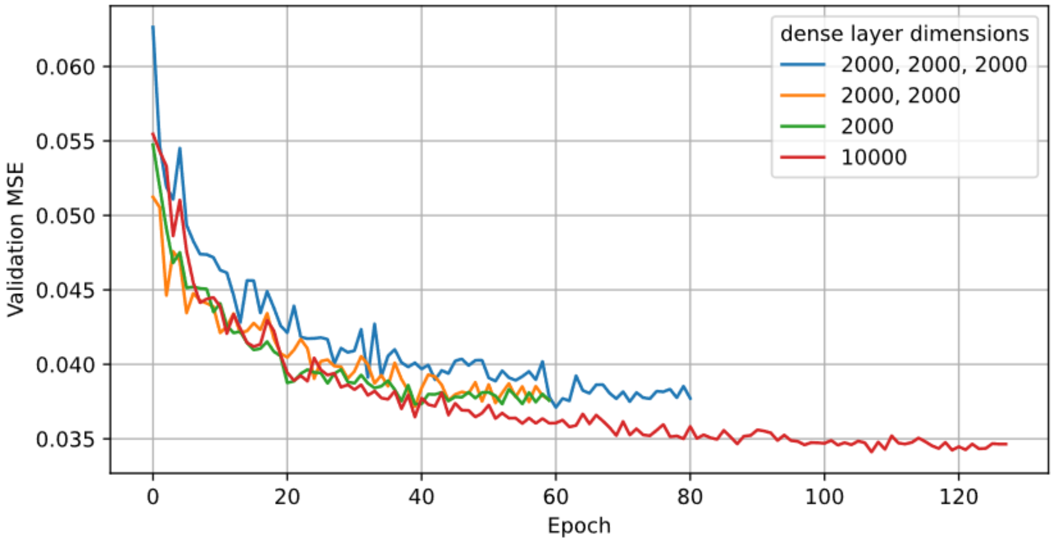


***Supplementary Figure 1.2: Models trained with varying model architectures.*** *The model architecture was varied to select a suitable model architecture. The models all used the same hyperparameters as the final model, except for the number of layers and dimensions of these layers. The input layer always consisted of 10000 bins with a width of 0.1 Da and a final embedding size of 500. The legend shows the dense layers used between the input layer and the predicted embedding, e.g., 2000, 2000, 2000 is a dense neural network with 3 fully connected layers, each with 2000 nodes. The validation MSE was calculated over the spectra in the validation test set. To calculate the validation MSE, the pairs were binned in 10 equal Tanimoto score bins between 0 and 1, the MSE was calculated per bin and the average was taken over the 10 bins.*

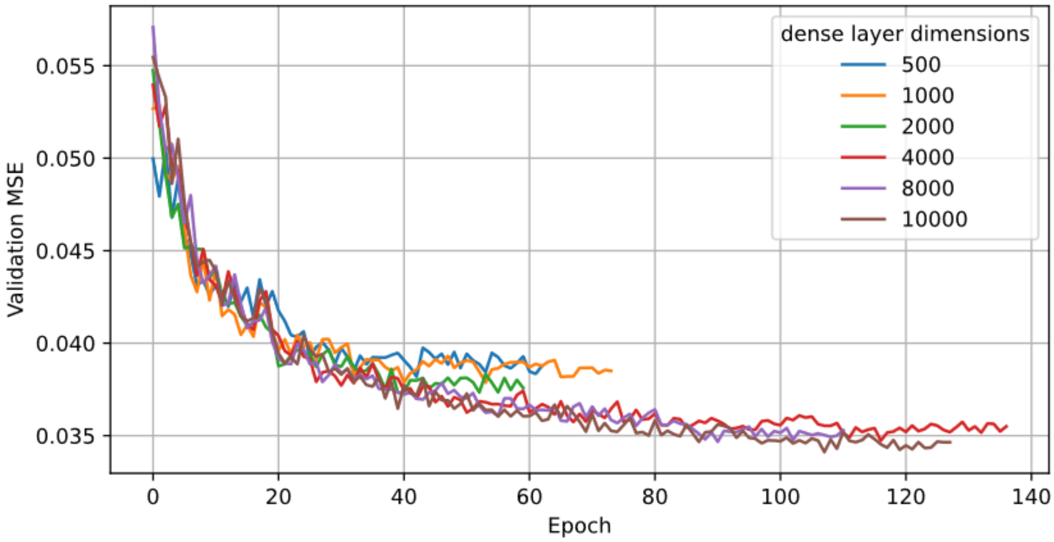


**Supplementary Figure 1.3: Models trained with a single dense neural network layer varying the dense layer dimension.** *The model architecture was varied to select a suitable model architecture.* The models all used a single dense layer between the input data and the predicted embedding. The dimension of this single layer was varied. *The validation MSE was calculated over the spectra in the validation test set.* To calculate the validation MSE, *the pairs were binned in 10 equal Tanimoto score bins between 0 and 1, the MSE was calculated per bin and the average was taken over the 10 bins.*

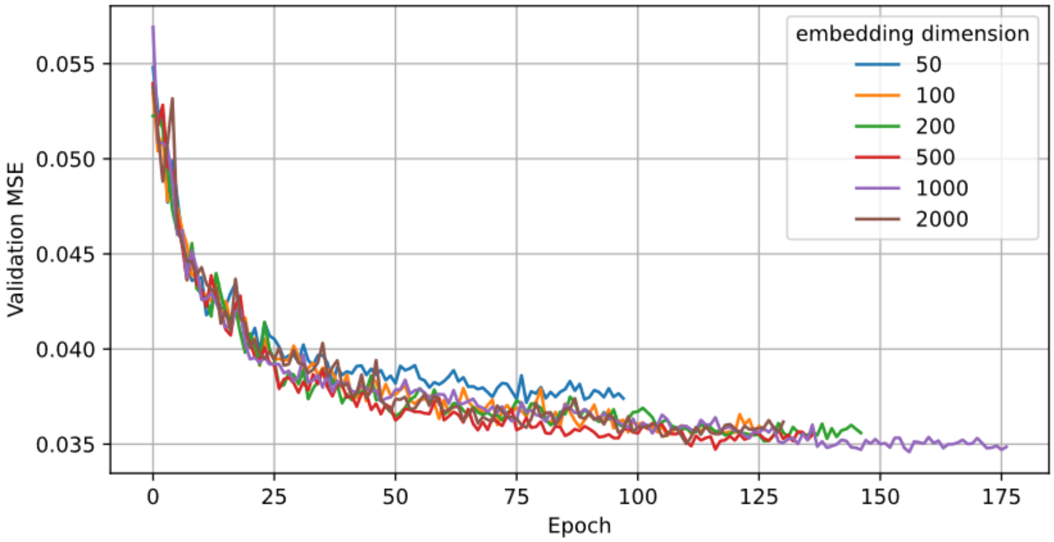


***Supplementary Figure 1.4: Average MSE per bin for different embedding dimensions.*** *The models all used the same hyperparameter settings, except for the embedding size used. The validation MSE was calculated over the spectra in the validation test set. The pairs were binned in 10 equal Tanimoto score bins between 0 and 1, the MSE was calculated per bin and the average was taken over the 10 bins.*

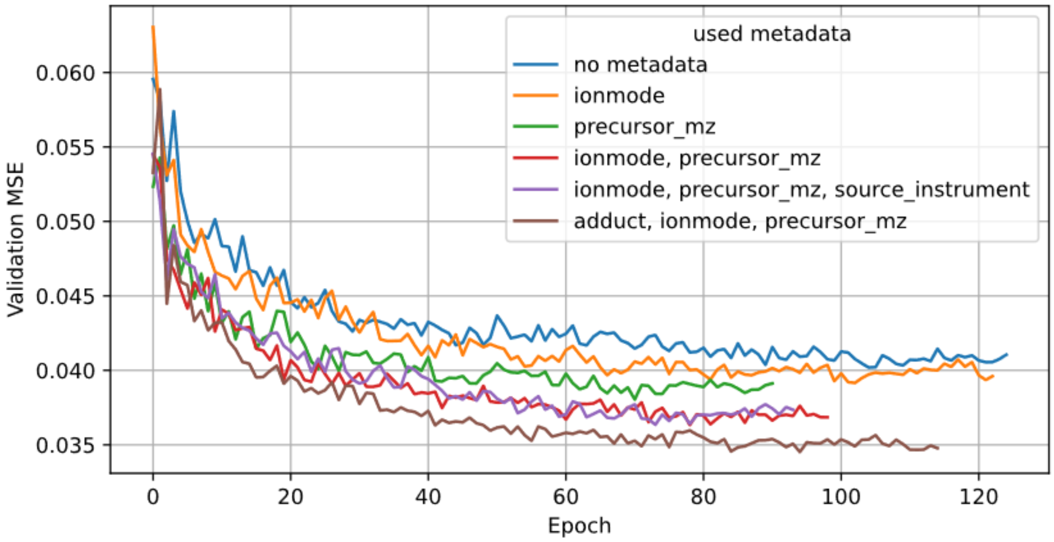


***Supplementary Figure 1.5: Average MSE per bin for dual-ionization mode model trained with different metadata input.*** *The models all used the same hyperparameter settings, except for the additional metadata used. An embedding size of 500 and a single layer of size 2000 were used. The validation MSE was calculated over the spectra in the validation test set. The pairs were binned in 10 equal Tanimoto score bins between 0 and 1, the MSE was calculated per bin and the average was taken over the 10 bins.*

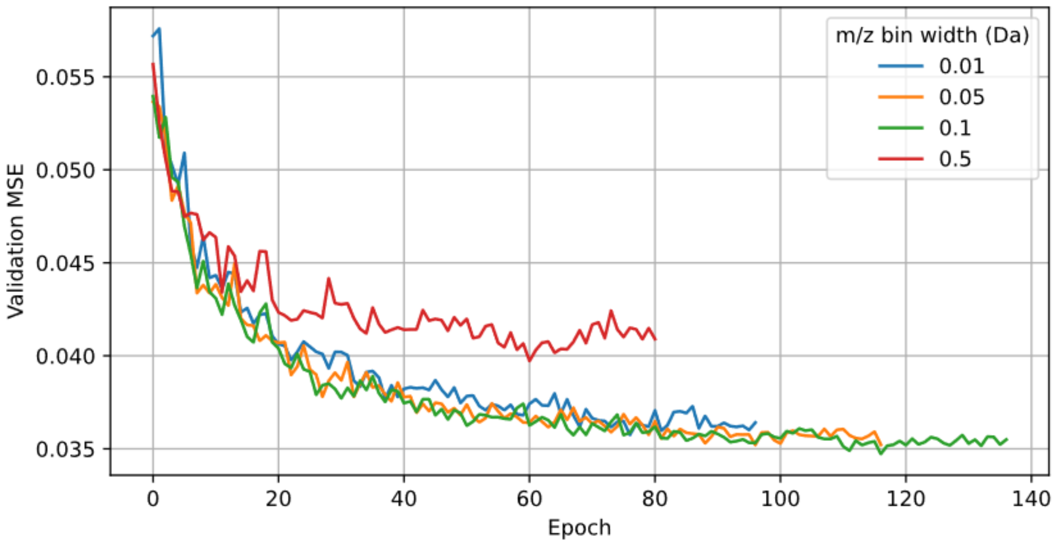


***Supplementary Figure 1.6: Average MSE per bin for dual-ionization mode model trained with different m/z bin widths.*** *The models all used the same hyperparameter settings, except for the loss function used. The validation MSE was calculated over the spectra in the validation test set. The pairs were binned in 10 equal Tanimoto score bins between 0 and 1, the MSE was calculated per bin and the average is taken over the 10 bins.*

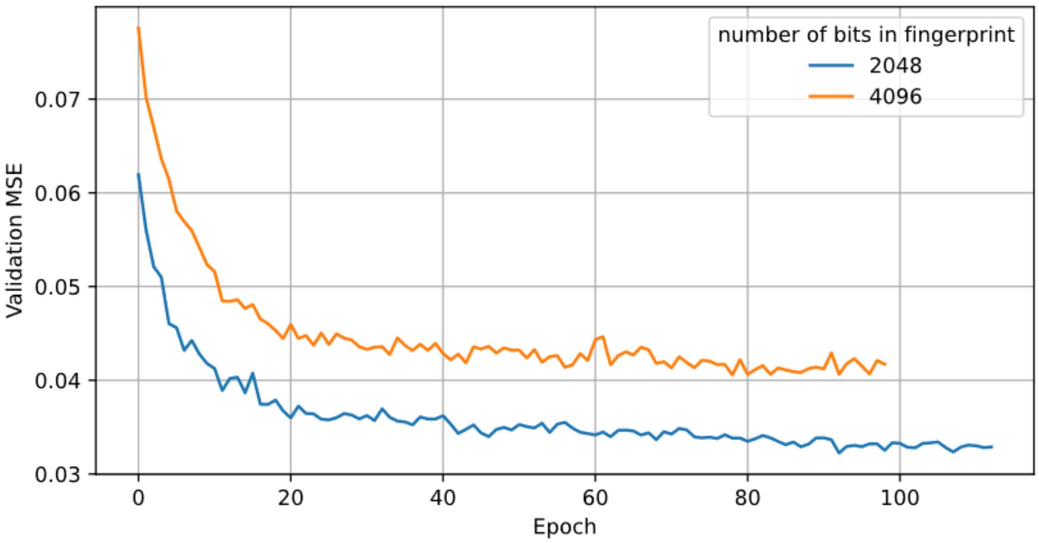


***Supplementary Figure 1.7: Average MSE per bin model trained to predict similarity based on fingerprints with different numbers of bits.*** *The models both used the same hyperparameter settings, except for the number of bits used for the Daylight fingerprint. The validation MSE was calculated over the spectra in the validation test set. The pairs were binned in 10 equal Tanimoto score bins between 0 and 1, the MSE was calculated per bin, and the average was taken over the 10 bins.*

### Supplementary Section 2. Pair sampling optimization

Sampling molecule pairs is a crucial step in optimizing model training. Sampling random pairs would result in a very large fraction of pairs having low similarity, since most molecule pairs have a low similarity score. To ensure a more equal sampling distribution, we sample equally from 10 equally spaced bins between 0 and 1. In all the sampling algorithm optimization tests in this section, except Supplementary Figure 2.6, we used settings resulting in sampling each molecule on average 100 times.

#### Balanced molecule sampling

Randomly sampling an equal number of pairs from each Tanimoto bin is possible, but results in unequal sampling of molecules, see Supplementary Figure 2.1a. This is not an efficient way of using the diversity in the training data and might result in inferior generalization.

In our sampling algorithm, we counteract this by tracking the molecule sampling frequency. During sampling, the least sampled molecule is picked, followed by sampling the least frequently sampled molecule that has a pair in this bin. This still results in some disbalance in molecule frequency, since for the second molecule of a pair there is sometimes only one or a few already frequently sampled options. To counteract this, we have implemented a maximum sampling frequency. If the maximum sampling frequency is reached the molecule will not be sampled even if it is the only available pair for the least sampled molecule in that bin. To enable setting a maximum molecule sampling frequency that is close to the average molecule frequency we need to allow resampling of pairs. Otherwise, it may result in an insufficient number of molecule pairs available to sample equally in each bin. Combined this results in both balanced sampling across Tanimoto bins and little variation in molecule sampling frequency.

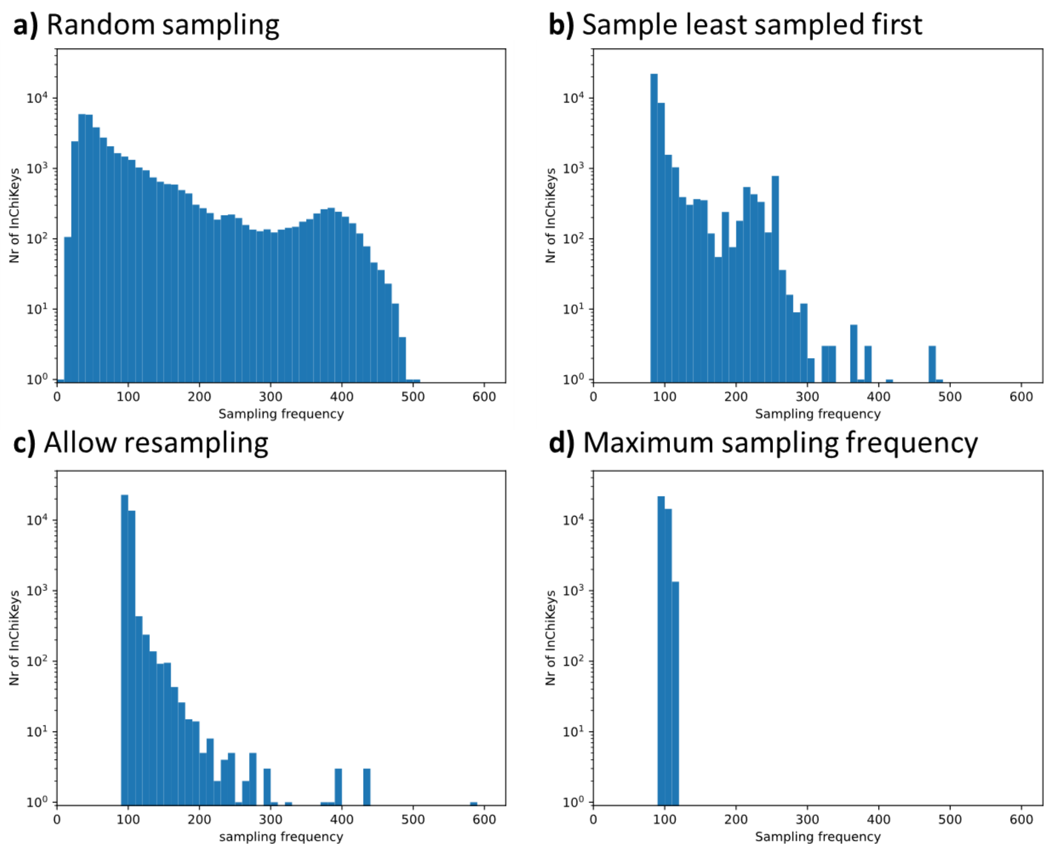


***Supplementary Figure 2.1: The molecule sampling frequency for different sampling algorithms.*** *The different sampling algorithms are all tested in the training set. The sampling frequency is the number of times a unique InChIKey is sampled.* ***a)*** *Random sampled an equal number of pairs from each bin.* ***b)*** *Sampled least frequently sampled molecules first.* ***c)*** *Sampled least frequently sampled molecules first, but allowed resampling of pairs that have already been sampled.* ***d)*** *Sampled least frequent molecules first, allowed resampling of pairs and set a maximum of molecule frequency at 110.*

#### Balanced score distribution per molecule

The sampling algorithm mentioned above results in perfect balance over the Tanimoto bins and almost equal sampling per molecule. However, per molecule the pair distribution is not balanced. For some pairs, mainly pairs with low Tanimoto scores are sampled, and for others mainly pairs with high Tanimoto scores are sampled. This imbalance can be due to biases in our training data, as some chemical classes might be more common in our dataset, while others might be unique molecules without similarity in this training set. It is not recommended to train a model that is able to use these potential biases during training. The risk is that the model learns that a molecule always has a low similarity during training. A model that uses these biases will not generalize well when predicting a highly chemically similar pair during deployment.

Ideally, each molecule would be sampled equally from each Tanimoto score bin. However, this is not possible, since many molecules have no pair available in some of the Tanimoto bins. These cases are unequally distributed over the bins, see Supplementary Figure 2.2. All molecules have at least one pair in the bin between 0.9-1.0, since all molecules have a Tanimoto score of 1, when comparing to itself.

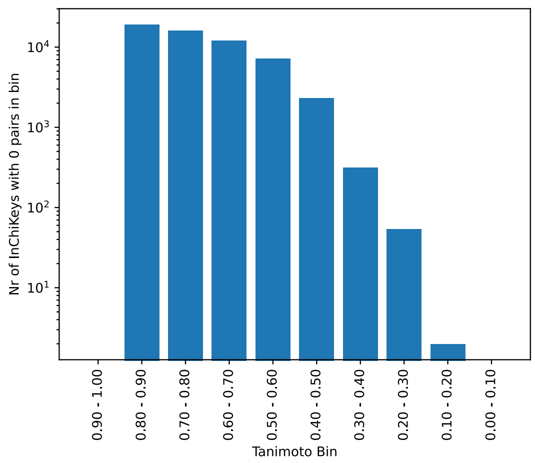


***Supplementary Figure 2.2: Molecule frequency of molecules with 0 pairs in each bin.*** *The number of unique InChIKeys with 0 pairs in each Tanimoto bin for the training spectra.*

#### Optimize bin order

The pairs are sampled per bin, molecules that are under sampled, since they don't have any pairs in a bin can be compensated in later bins. To make sure that every pair can actually be compensated by bins sampled later, the best approach is to start in the bin with the largest number of molecules with 0 pairs. This leaves more options for resampling in the following bins. However, most of the missing molecules in the bins 0.6-0.9 would in that case be oversampled in the bins 0.6-0.4, resulting in these molecules having a low average Tanimoto score. Ideally molecules with no pairs in bins are compensated in bins with similar Tanimoto scores, e.g. compensating missing pairs between 0.8-0.9 in 0.9-1.0 or 0.7-0.8. To detect which bin order achieves this best, the sampling algorithm was run multiple time, while varying the position of the 0.9-1.0 bin, while the other bins were kept constant from high to low. The average Tanimoto score per molecule was calculated and the fraction of pairs in the most sampled bin per molecule was calculated. These were used as a metric for testing how well the pairs were distributed over the Tanimoto score bins per InChiKey. Based on Supplementary Figure 2.3, we decided to use the bin order where we sample the bin between 0.9 and 1.0 as the third bin. This results in the average score per molecule shown in Supplementary Figure 2.4b.

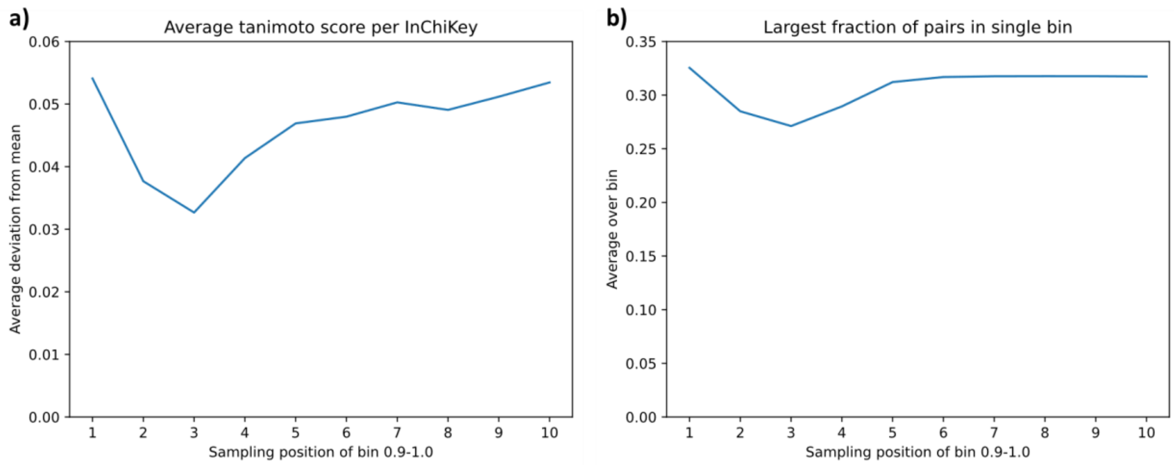


***Supplementary Figure 2.3: Effect of sampling order.*** *The sampling algorithms samples pairs per bin, in later bins under sampled InChIKeys can be compensated. The sampling order has an effect on how this oversampling happens. To test the optimal sampling order, the sampling order was varied. The position of the 0.9-1.0 bin was varied, the other bins are ordered from high to low. The effect is tested on the training spectra.* ***a)*** *For each molecule the average of the scores for the selected pair was calculated. The absolute difference with 0.5 (the mean) was calculated, followed by calculating the average over all molecules.* ***b)*** *For each molecule the distribution of pairs over the bins was calculated. The bin with the highest number of pairs was selected and the fraction of the total number of pairs was calculated for this molecule. A perfectly balanced score would result in 0.1 as average per bin (total/ number of bins).*

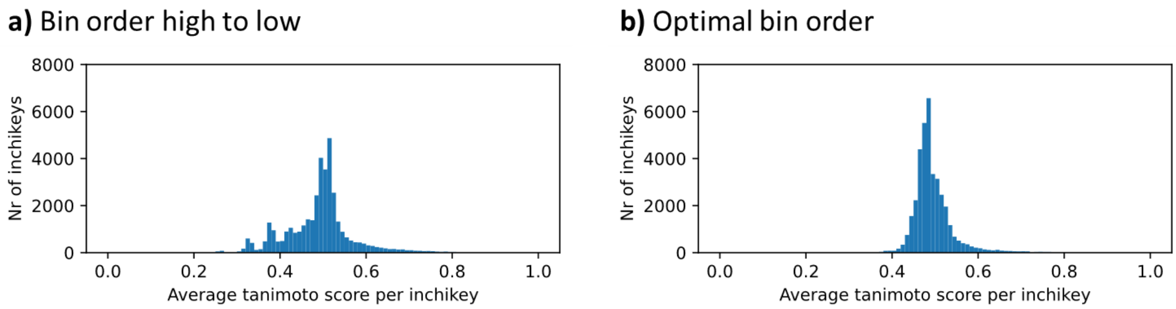


***Supplementary Figure 2.4: Average Tanimoto score over pairs sampled per molecule for the training spectra.*** ***a)*** *Sampling with bin order from high to low.* ***b)*** *Sampling with bin order [(0.8, 0.9), (0.7, 0.8), (0.9, 1.0), (0.6, 0.7), (0.5, 0.6), (0.4, 0.5), (0.3, 0.4), (0.2, 0.3), (0.1, 0.2), (-0.01, 0.1), ]*

#### Resampling

To achieve a well-balanced Tanimoto score distribution, it is required to allow some resampling of already sampled molecule pairs. To minimize the resampling of pairs we keep track of the sampling frequency of each pair. Before selecting the least frequently sampled second molecule, we select the least frequently sampled pairs. If no limit is set, this still results in resampling some pairs 50 times, see Supplementary Figure 2.5a. Implementing a maximum resampling rate can reduce the frequency of resampling molecule pairs. Supplementary Figure 2.5 shows that setting a low maximum resampling rate results in a bad score distribution per molecule, while the number of unique pairs selected is barely affected by the resampling rate. Based on these figures, a maximum pair resampling of 20 seemed optimal. However, we recommend not to set a maximum resampling value, since the effect is minimal, and setting a too low maximum resampling setting can result in a bad balance of pairs selected per molecule.

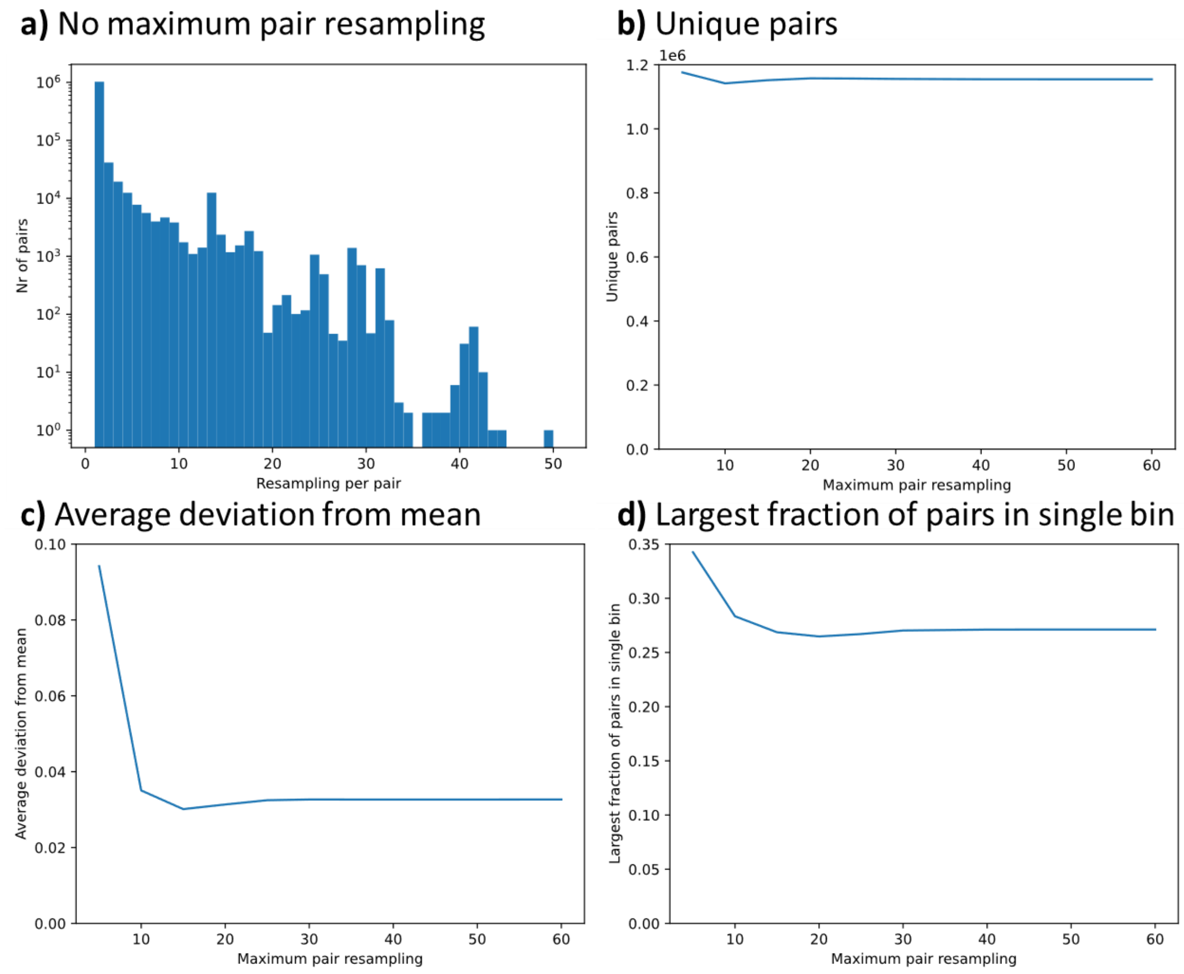


***Supplementary Figure 2.5: Effect of setting a maximum resampling on the training spectra. a)*** *The frequency of sampling each molecule pair without maximum resampling rate.* ***b)*** *The number of unique pairs sampled for different maximum pair resampling rates.* ***c)*** *For different maximum pair resampling rates the deviation from the mean Tanimoto score per molecule was calculated. For each molecule the average of the scores for the selected pair was calculated. The absolute difference with 0.5 (the mean) is calculated, followed by calculating the average over all molecules.* ***d)*** *For different maximum pair resampling rates the deviation from the mean Tanimoto score per molecule was calculated. For each molecule the distribution of pairs over the bins was calculated. The bin with the highest number of pairs was selected and the fraction of the total number of pairs was calculated for this molecule. A perfectly balanced score would result in 0.1 as average per bin (total/ number of bins).*

#### Number of sampled pairs

The number of pairs sampled can be varied. This can be changed by changing the average sampling count per molecule. By increasing the number of sampled pairs, the unique number of pairs increases most in the lower bins with many available pairs, while in the higher bins, with less available pairs, the number of unique pairs increases slower, since the number of pairs is mostly increased by resampling the previously selected pairs more frequently, see Supplementary Figure 2.6. Increasing the number of pairs sampled increases the runtime of the sampling algorithm. Sampling each molecule more than 100 times did not improve model performance (see Supplementary Figure 2.6b), therefore we settled for sampling each molecule 100 times on average.

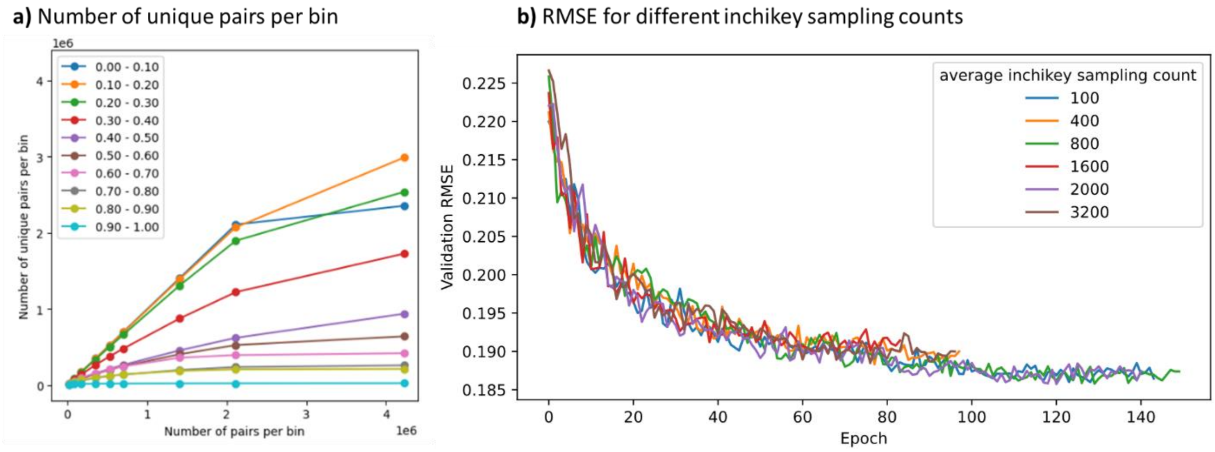


**Supplementary Figure 2.6: Effect of number of pairs sampled. a)** Number of unique pairs per bin for different numbers of pairs sampled from the training spectra. **b)** The validation RMSE for models trained for different average InChIKey sampling counts. This shows the number of pairs sampled per InChiKey does not have a large effect on training.

### Supplementary Section 3. Comparison to MS2DeepScore 0.2.0 model

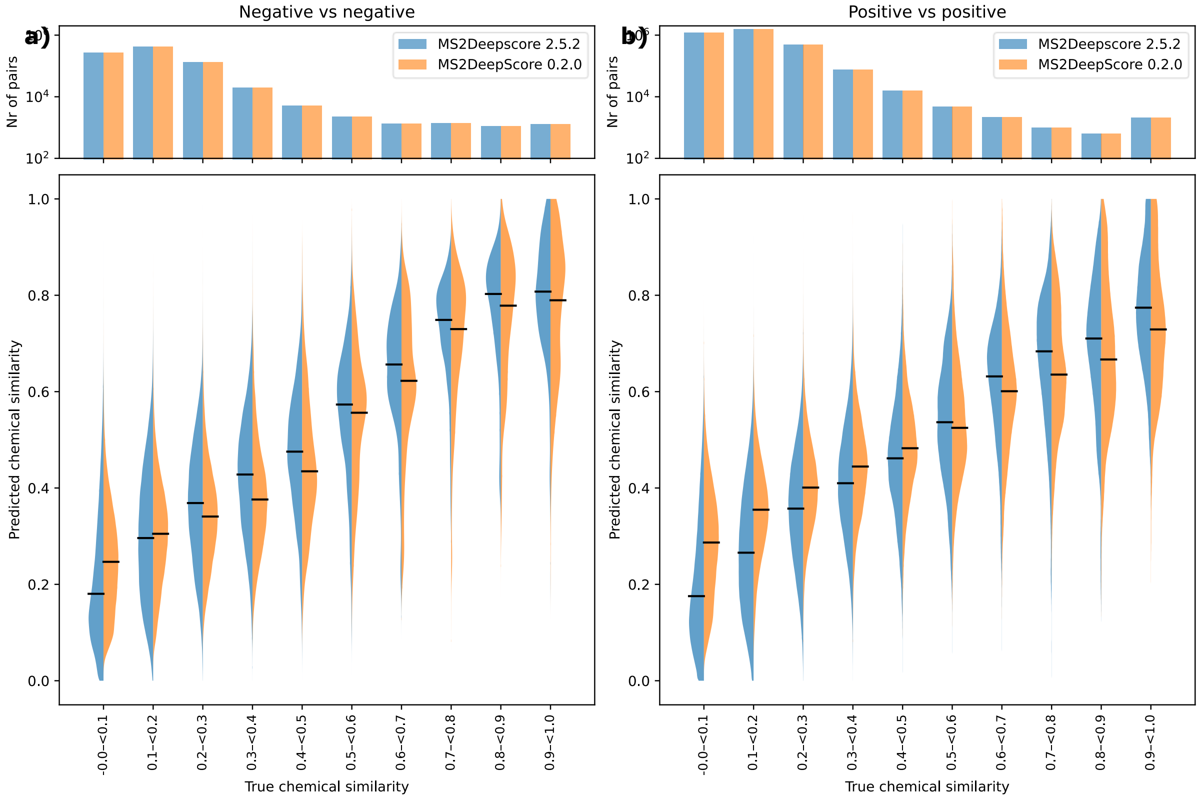


***Supplementary Figure 3.1: Comparison between current MS2DeepScore version (2.5.2) and the original MS2DeepScore version (0.2.0).*** *The MS2DeepScore model 0.2.0 is the version used to train the model with the same architecture as the original MS2DeepScore paper. This model is retrained on the same training data and benchmarked on the same test set to enable comparison to the new MS2DeepScore version (2.5.4). For the 0.2.0 architecture, two models are trained, one on positive ionization mode and one on negative ionization mode spectra. Predictions are made between all test spectra, followed by taking the average per unique molecule pair. The violin plots show the kernel density estimation (KDE) of the predicted values, the black lines represent the median. The bar plot on the top shows the log-scaled count of the number of unique molecule pairs in each bin with the corresponding chemical similarity. The metric used for chemical similarity prediction is the Tanimoto score between Daylight fingerprints.* ***a)*** *Predictions between pairs of negative ionization mode spectra.* ***b)*** *Predictions between pairs of positive ionization mode spectra.*

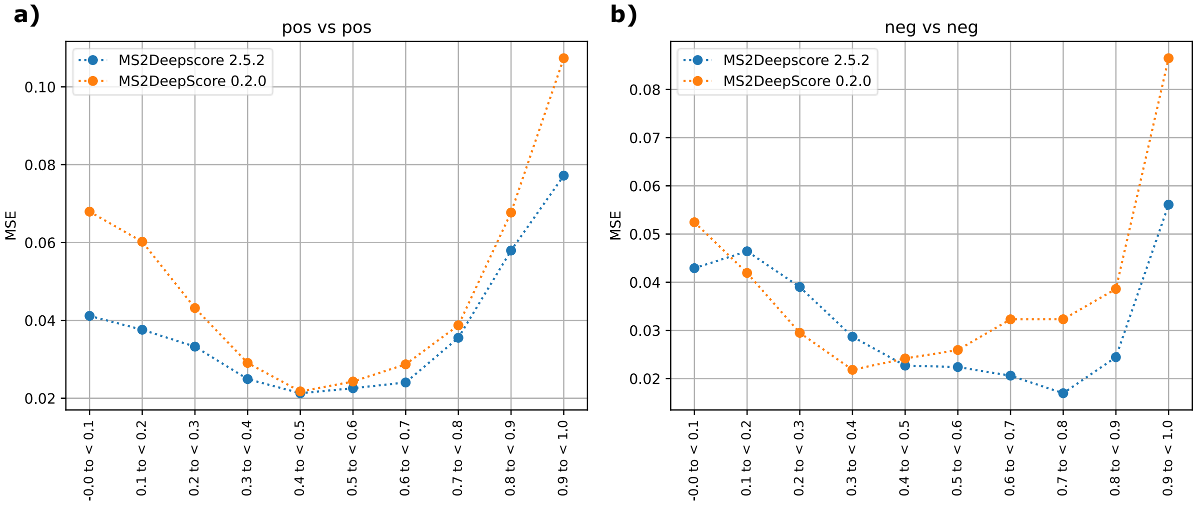


**Supplementary Figure 3.2: MSE per Tanimoto bin for current MS2DeepScore version (2.5.4) versus original MS2DeepScore version (0.2.0).** *Compound pairs are sampled from the test set per Tanimoto bin. The average MSE per compound pair is calculated followed by calculating the average over all compound pairs in the Tanimoto bin.* ***a)*** *Predictions between pairs of positive ionization mode spectra.* ***b)*** *Predictions between pairs of negative ionization mode spectra.*

### Supplementary Section 4: Comparison to model trained on single ionization mode

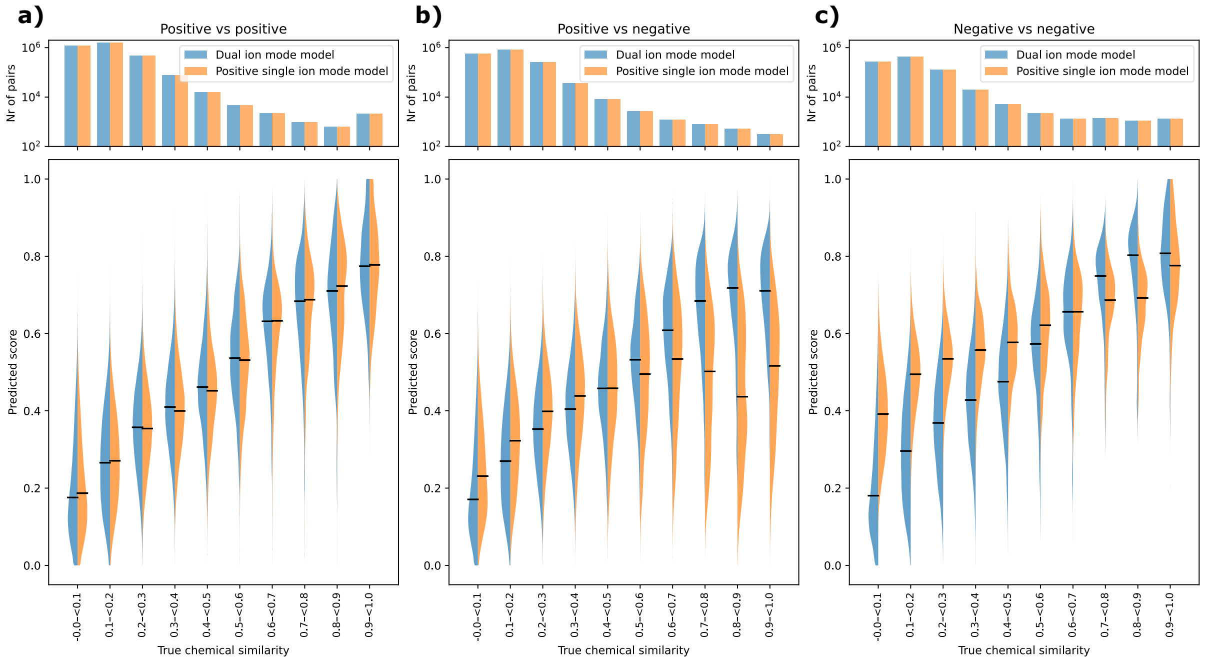


**Supplementary Figure 4.1*****: Comparison between cross-ionization model and model trained only on positive ionization mode spectra.*** *The model trained on only positive ionization mode spectra used the same architecture, but only trained on the positive ionization mode spectra. Predictions are made between all test spectra, followed by taking the average per unique molecule pair. The violin plots show the kernel density estimation (KDE) of the predicted values, the black lines represent the median. The bar plot on the top shows the log-scaled count of the number of unique molecule pairs in each bin with the corresponding chemical similarity. The metric used for chemical similarity prediction is the Tanimoto score between Daylight fingerprints.* ***a)*** *Predictions between pairs of positive ionization mode spectra.* ***b)*** *Predictions between pairs of positive and negative ionization mode spectra.* ***c)*** *Predictions between pairs of negative ionization mode spectra.*

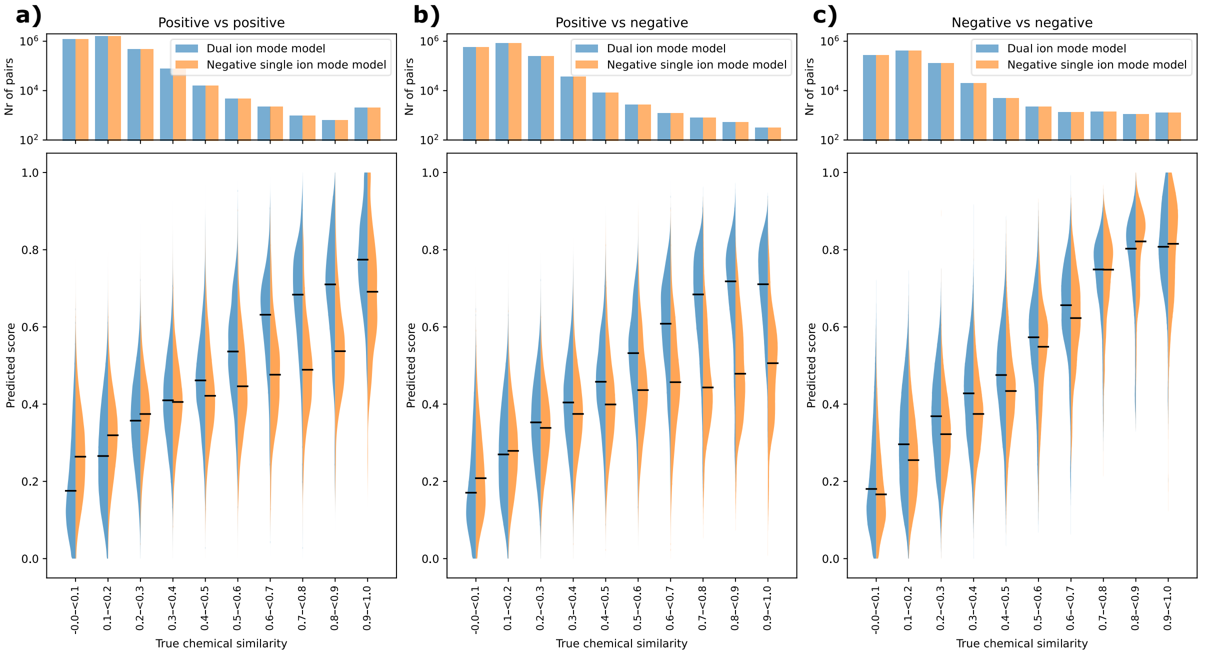


**Supplementary Figure 4.2: *Comparison between cross-ionization model and model trained only on negative ionization mode spectra.*** *The model trained on only negative ionization mode spectra used the same architecture, but only trained on the negative ionization mode spectra. Predictions are made between all test spectra, followed by taking the average per unique molecule pair. The violin plots show the kernel density estimation (KDE) of the predicted values, the black lines represent the median. The bar plot on the top shows the log-scaled count of the number of unique molecule pairs in each bin with the corresponding chemical similarity. The metric used for chemical similarity prediction is the Tanimoto score between Daylight fingerprints.* ***a)*** *Predictions between pairs of positive ionization mode spectra.* ***b)*** *Predictions between pairs of positive and negative ionization mode spectra.* ***c)*** *Predictions between pairs of negative ionization mode spectra.*

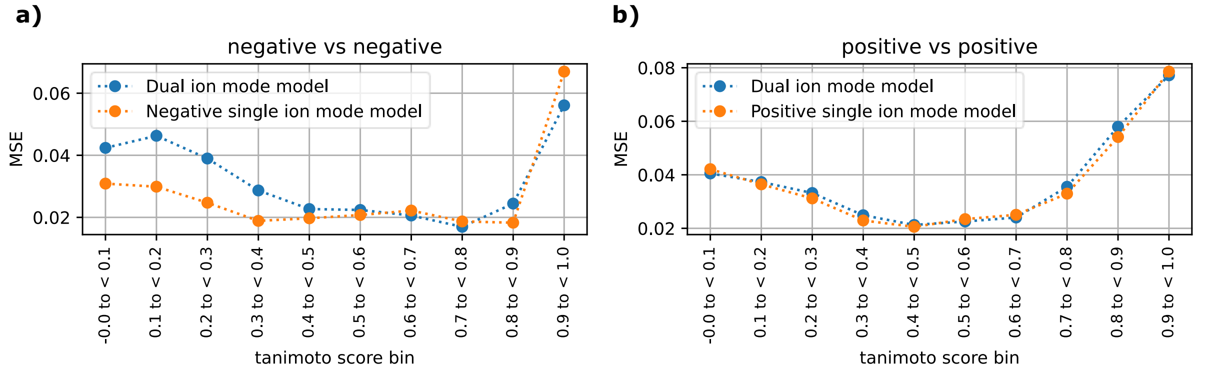


**Supplementary Figure 4.3: MSE per Tanimoto bin for single ionization mode model compared to dual ionization mode model.** *Compound pairs are sampled from the test set per Tanimoto bin. The average MSE per compound pair is calculated followed by calculating the average over all compound pairs in the Tanimoto bin.* ***a)*** *Predictions between pairs of negative ionization mode spectra. The model trained on single ionization mode spectra performs better than the model trained on both ionization modes.* ***b)*** *Predictions between pairs of positive ionization mode spectra.*

### Supplementary Section 5. Comparison to modified cosine score

The modified cosine score is not designed for predicting Tanimoto scores, so it is not expected to correlate well with the Tanimoto score. The strength of the modified cosine score is that when a high score is predicted it most of the time is a high similarity match, however, for many highly similar metabolites the modified cosine score is 0. Since the compute time of modified cosine score is significantly higher than for MS2DeepScore, it was not feasible to calculate an all vs all prediction matrix for the test set. Instead 1 spectrum per unique molecule was selected from the test set resulting in 1830 positive ionization mode spectra and 924 negative ionization mode spectra. Instead of 24911 positive mode spectra and 7142 negative mode spectra. In addition the spectra were first filtered to contain only the 100 highest intensity peaks.

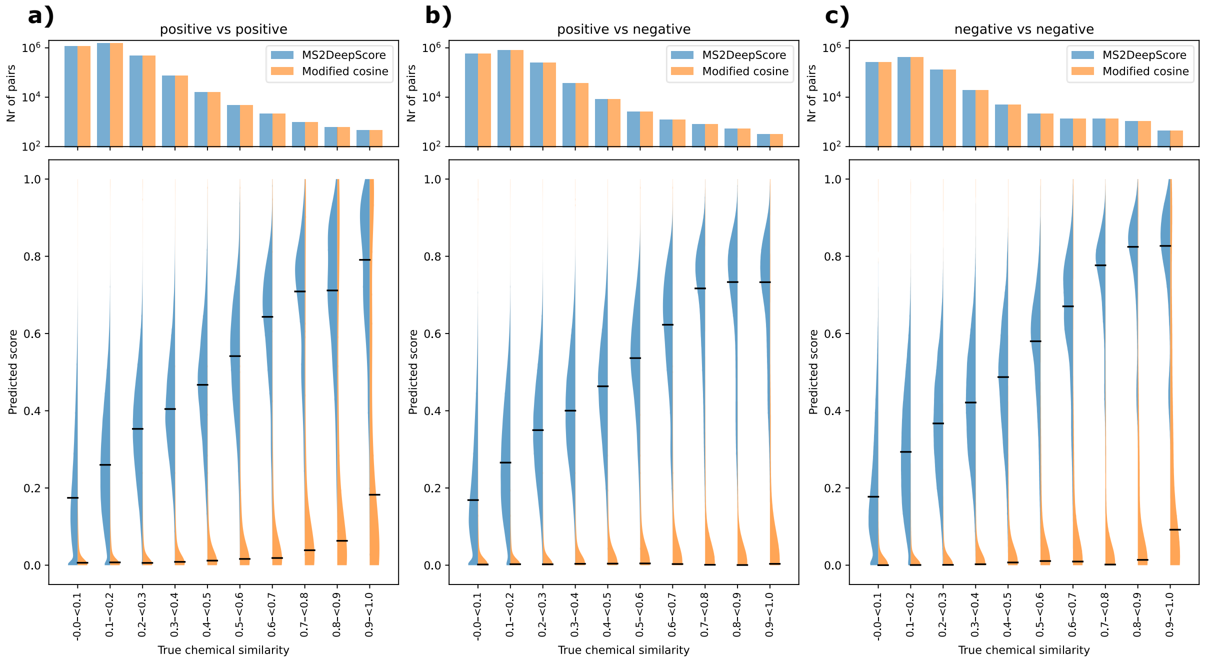


***Supplementary Figure 5.1: Side by side violin plot of modified cosine score predictions and MS2DeepScore predictions.*** *One spectrum per unique molecule is randomly sampled from the test set, resulting in a subset of the test set. Modified cosine score predictions and MS2DeepScore predictions are made between all sampled spectra. The violin plots show the kernel density estimation (KDE) of the predicted values, the black line represents the median for each bin. The bar plot shows the number of unique molecule pairs for each true chemical similarity bin. The true chemical similarity used is the Tanimoto score between Daylight fingerprints.* ***a)*** *Predictions between pairs of positive ionization mode spectra.* ***b)*** *Predictions between pairs positive and negative ionization mode spectra. Interestingly, MS2DeepScore models trained on only one of the ionization modes show a better than random prediction performance when predicting mass spectral similarity of spectra obtained in the other ionization mode or cross-ionization mode. This suggests that MS2DeepScore is able to detect patterns that generalize between the two ionization modes.* ***c)*** *Predictions between pairs of negative ionization mode spectra.*

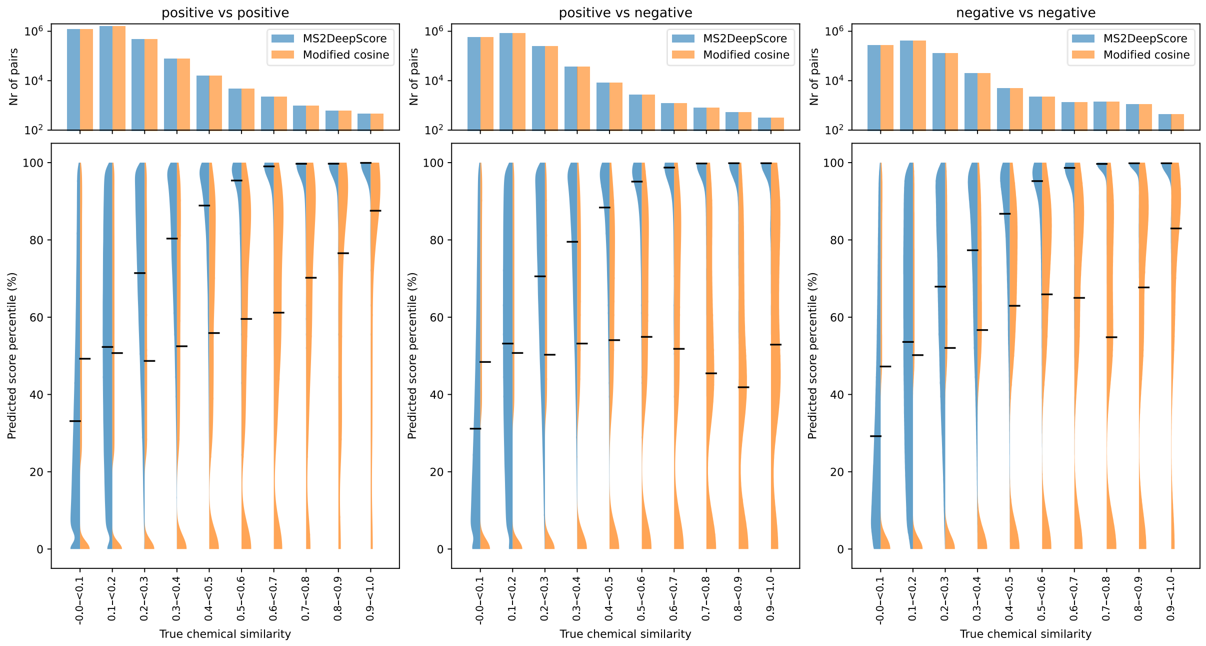


***Supplementary Figure 5.2: Side by side violin plot of modified cosine score prediction percentiles and MS2DeepScore prediction percentiles.*** *One spectrum per unique molecule is randomly sampled from the test set, resulting in a subset of the test set. Modified cosine score predictions and MS2DeepScore predictions are made between all sampled spectra. The predictions are converted to predicted percentiles, by ranking the scores over all predictions made for the used small test set of each ion mode, this enables a better comparison, since it converts the predictions to a more comparable scale. The violin plots show the kernel density estimation (KDE) of the predicted values, the black line represents the median for each bin. The bar plot shows the number of unique molecule pairs for each true chemical similarity bin. The true chemical similarity used is the Tanimoto score between Daylight fingerprints.* ***a)*** *Predictions between pairs of positive ionization mode spectra.* ***b)*** *Predictions between pairs positive and negative ionization mode spectra. Interestingly, MS2DeepScore models trained on only one of the ionization modes show a better than random prediction performance when predicting mass spectral similarity of spectra obtained in the other ionization mode or cross-ionization mode. This suggests that MS2DeepScore is able to detect patterns that generalize between the two ionization modes.* ***c)*** *Predictions between pairs of negative ionization mode spectra.*

### Supplementary Section 6. Embedding Evaluator

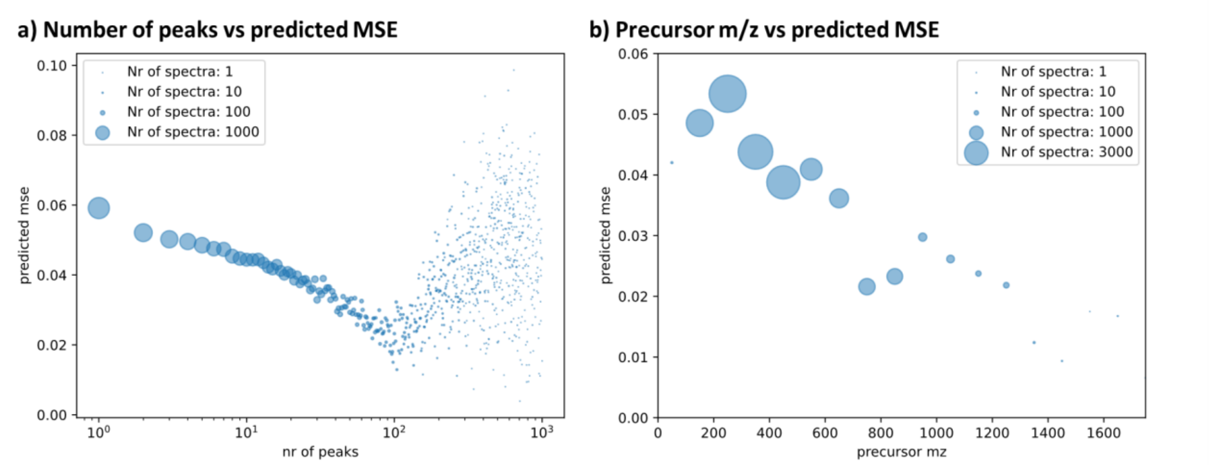


**Supplementary Figure 6.1: Relationship between predicted MSE and number of peaks and precursor *m/z*.** For all test spectra the MSE is predicted using our Embedding Evaluator model. **a)** The number of peaks is plotted against the predicted MSE. The average of all spectra with a specific number of peaks is plotted. The size of the dot shows the number of spectra in the test set that have this number of peaks. **b)** The precursor *m/z* is plotted against the predicted MSE. The average of all spectra in bins of 100 Da is plotted. The size of the dot shows the number of spectra in the test set that fall in this precursor *m/z* bin.

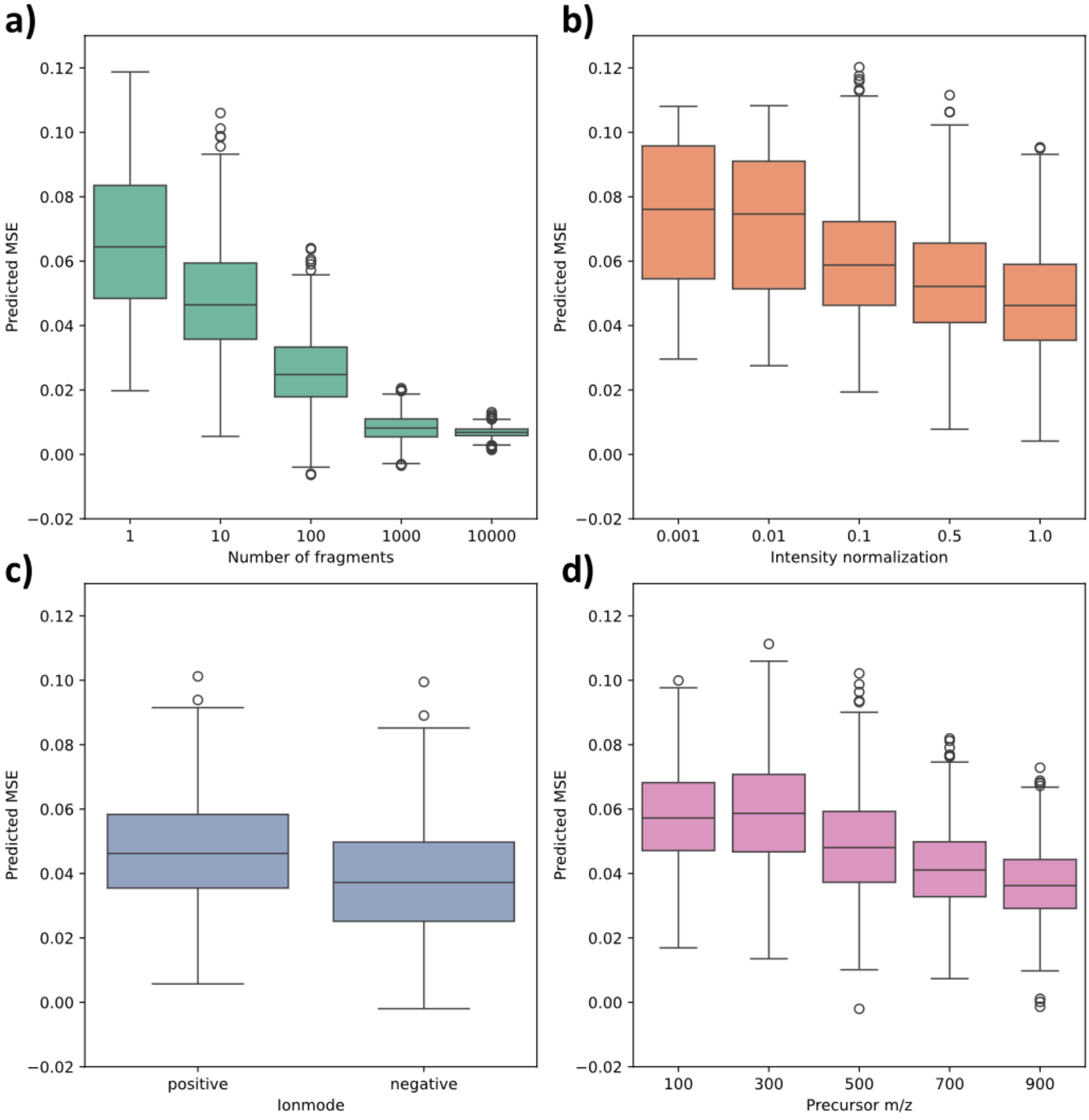


***Supplementary Figure 6.2: Predicted MSE for randomly generated spectra.*** *A 1000 spectra are randomly generated for each test by creating random fragments, with random intensities and random precursor m/z. The default number of fragments was 10, the default intensity was between 0 and 1, the default m/z values of the fragments were between 10 and 1000 Da, the default ionization mode was positive and a random precursor m/z was selected between 0 and 1000 Da.* ***a)*** *The number of fragments per randomly generated spectrum is varied.* ***b)*** *The maximum randomly generated intensity of the peaks is varied, e.g. 0.01 means random intensities are generated between 0 and 0.01.* ***c)*** *The ionization mode for all spectra is set to positive or negative.* ***d)*** *The precursor m/z is fixed to a specific value.*

### Supplementary Section 7. Detailed analysis of MS2DeepScore model

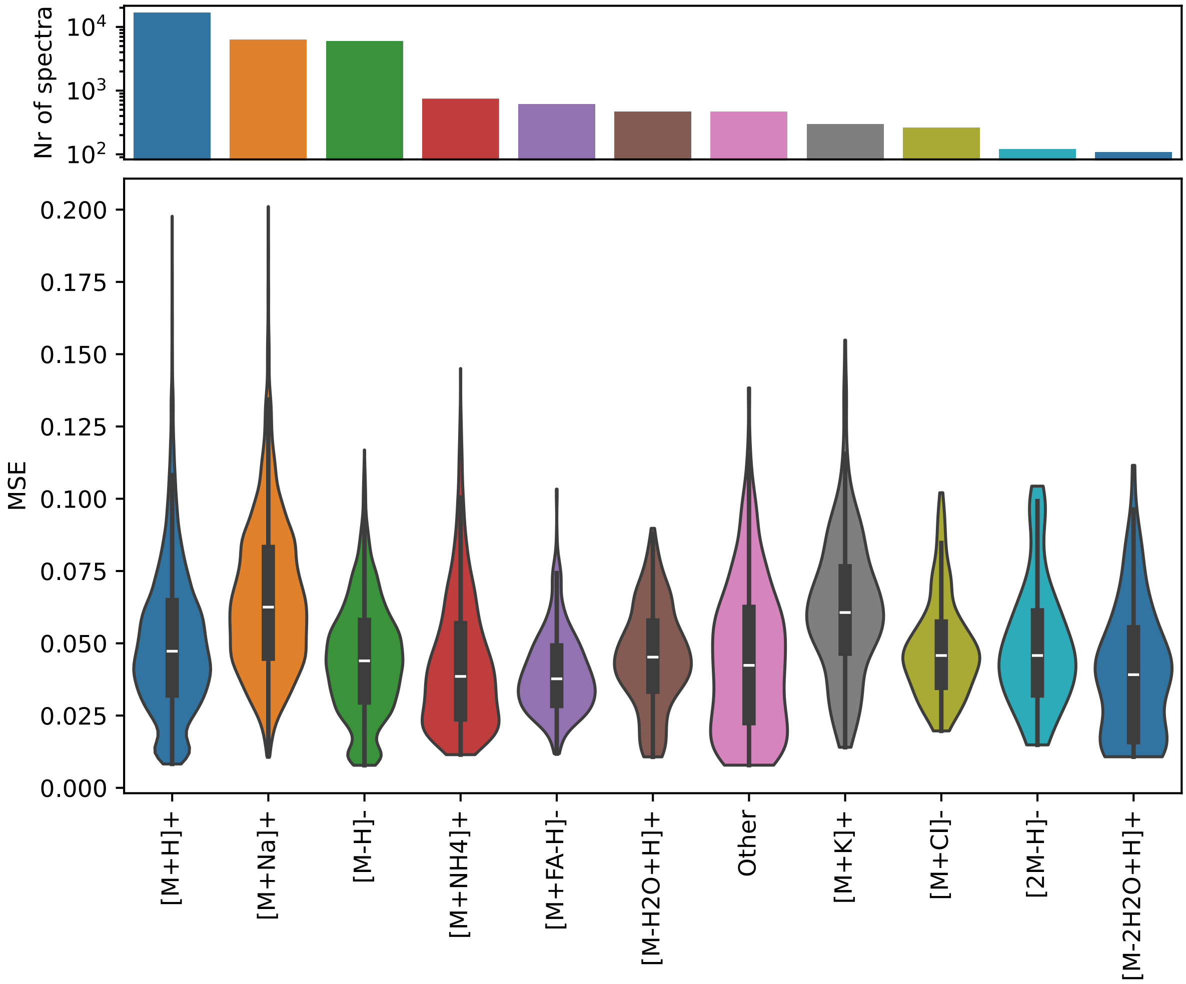


***Supplementary Figure 7.1: MSE per adduct type.*** *For the test set the MSE per spectrum was calculated. Here the distribution of MSE per adduct type is given. Any adduct with less than 100 spectra is combined in the "other" boxplot.*

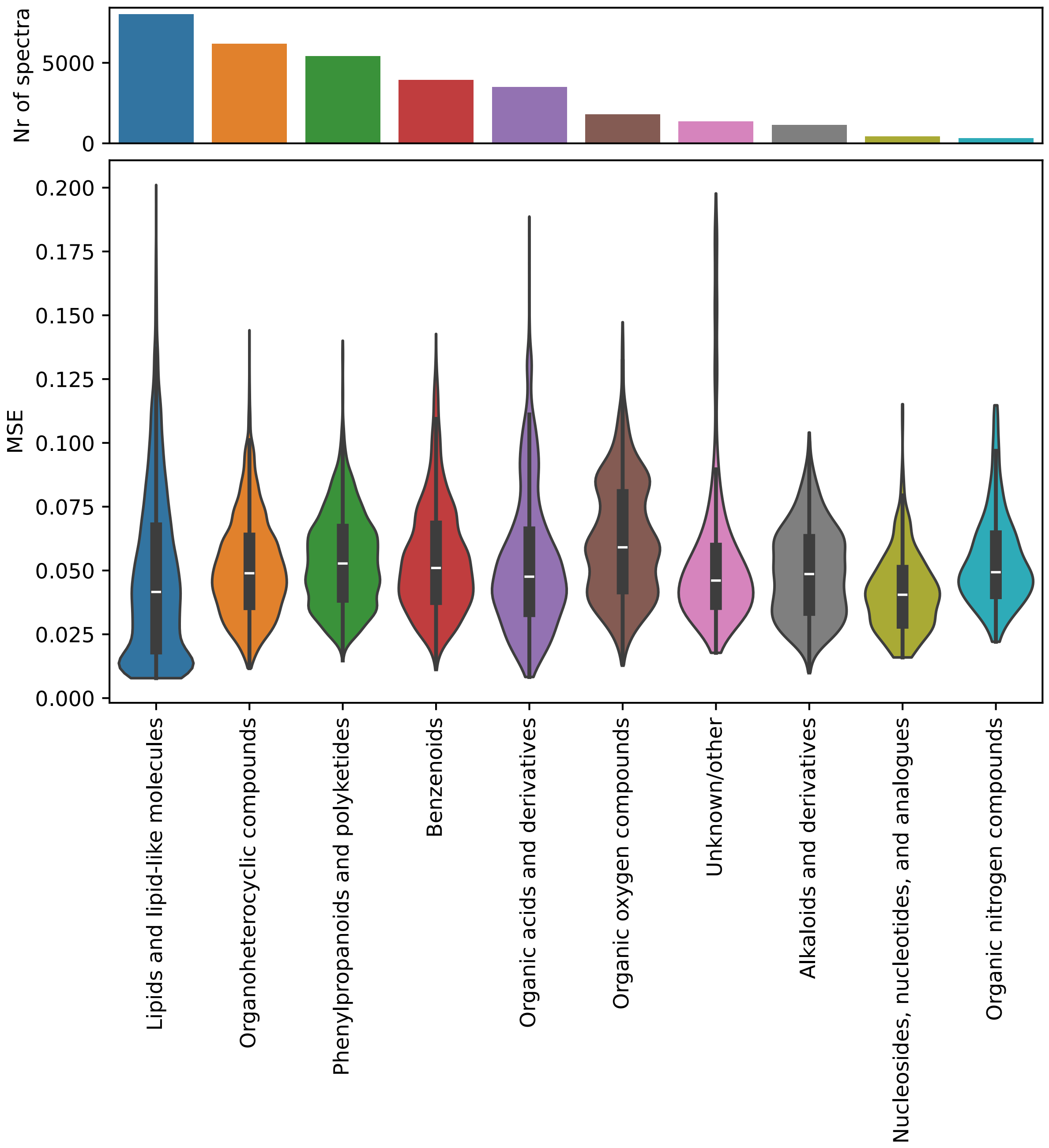


***Supplementary Figure 7.2: MSE per chemical class.*** *For the test set the MSE per spectrum was calculated. Here the distribution of MSE per chemical class is given. The chemical class is determined by using ClassyFire[1], both molecules for which no ClassyFire annotation was available and chemical classes with less than 300 spectra are combined in the boxplot Unknown/other.*

#### Supplementary Section 8. Case study mass spectrometer parameters

Mass spectrometric analysis of urine fractions plasma NIST SRM1950 was performed in positive and negative ionization modes (ESI+ and ESI-). The mass spectrometry parameters were set as follows for urine RP profiling: capillary voltage 1.5 kV (ESI+) and 1 kV (ESI-), sample cone voltage 20 V, source temperature 120°C, desolvation temperature 600°C, desolvation gas flow 1000 L/h, and cone gas flow 150 L/h. Data were collected in centroid mode with a scan range of 50-1200 *m/z* with a scan time of 0.1 s. For mass accuracy, LockSpray mass correction was performed using a 600 pg/µL leucine enkephalin solution (*m/z* 556.2771 in ESI+ and 554.2615 in ESI-) in 1:1 water:ACN solution at a flow rate of 15 µL/min. Lockmass scans were collected every 60 s and averaged over 4 scans. The mass spectrometer was operating in Fast DDA mode. The intensity threshold of the precursor ion was set to 100 K to trigger $MS^2$ fragmentation that was performed in centroid mode with a scan range of 50-1200 *m/z* and a scan time of 0.25 s. $MS^2$ was switched back to MS survey function after 2 s acquisition. Deisotoped peak selection option was enabled. The collision energy was set to the ramp of 15–30 eV and 30-60 eV for $MS^2$ acquisition of low and high mass ions, respectively.

#### Supplementary Section 9. Human blood plasma case study

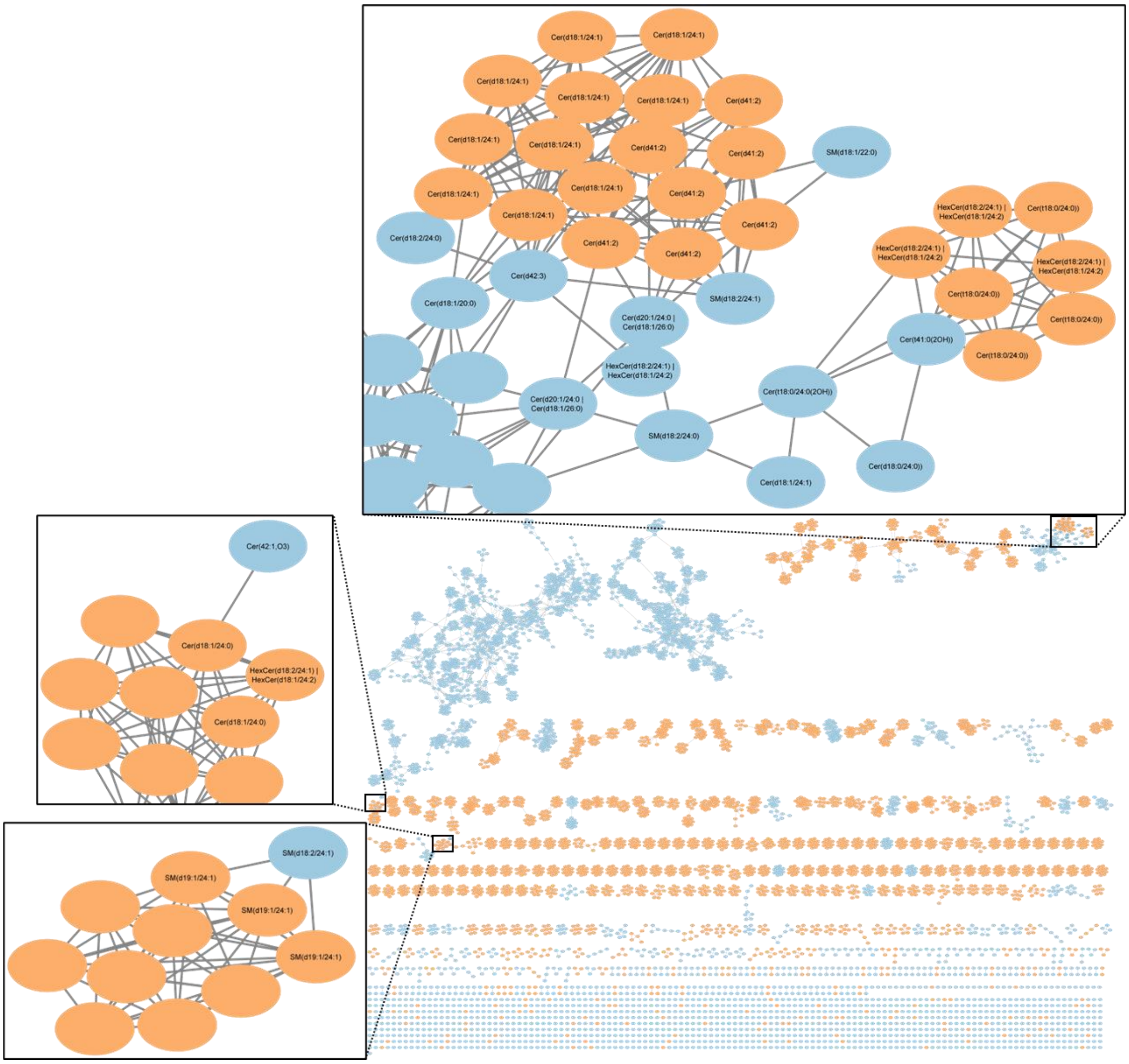


**Supplementary *Figure 9: Molecular network created with MS2DeepScore cross-ionization-mode model on human blood plasma case study.*** *By predicting chemical similarity between both the positive and negative ionization mode spectra, spectra of both ionization modes can be visualized together. An edge is created for an MS2DeepScore larger than 0.85. We highlight a few examples where MS2DeepScore was able to predict close chemical similarity between positive and negative ionization modes. The annotations were added by manual annotations.*

### Supplementary Section 10. Spectrum comparison plots

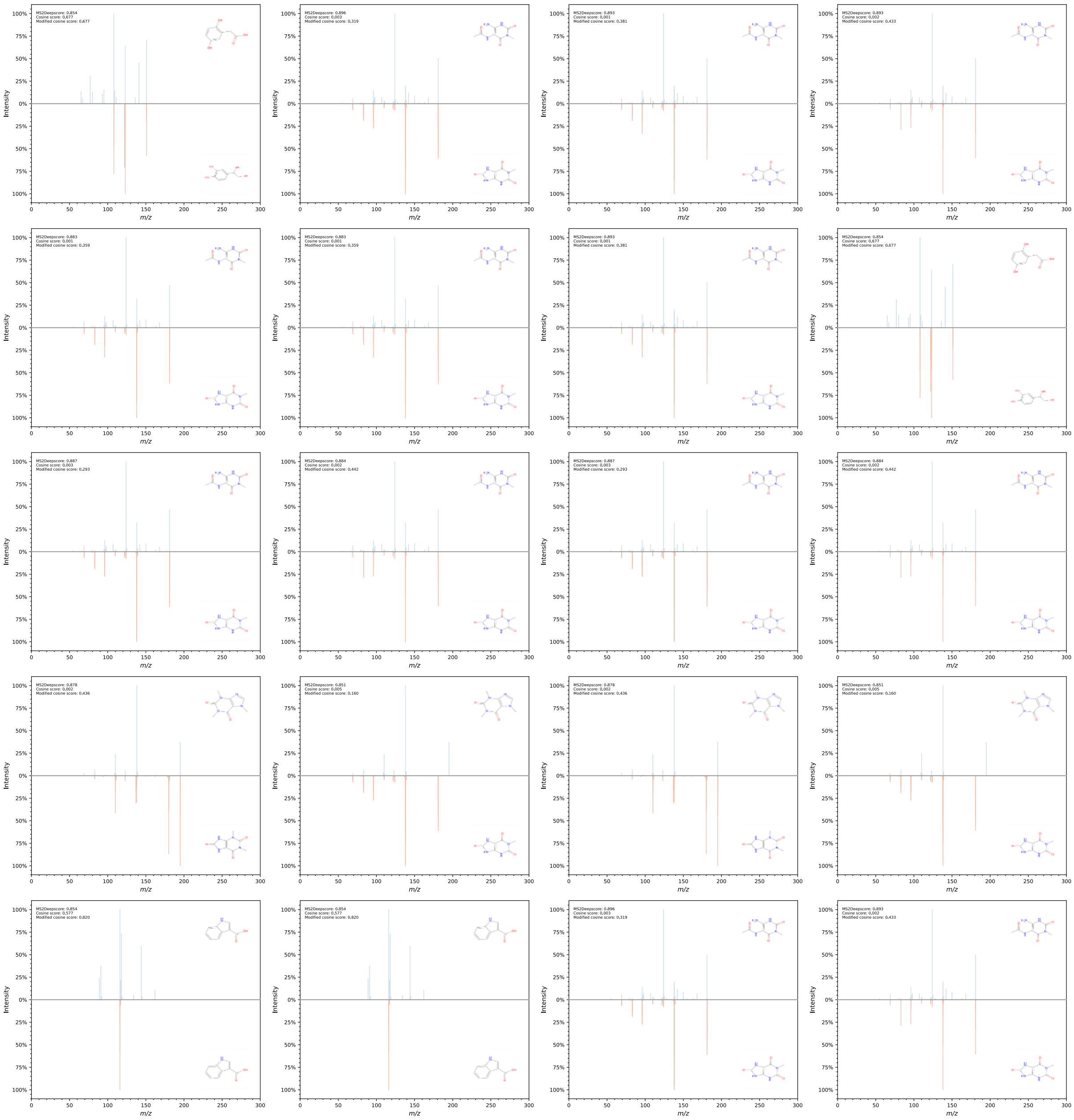

***Supplementary Figure 10.1: Spectrum comparisons of positive and negative ionization mode spectra for which MS2DeepScore predicts high chemical similarity.*** *The spectra are from the urine case study. Positive ionization mode spectra are blue and negative ionization mode spectra are orange. All annotated spectra connected across ionization modes in the molecular network in Figure 3 are visualized.*

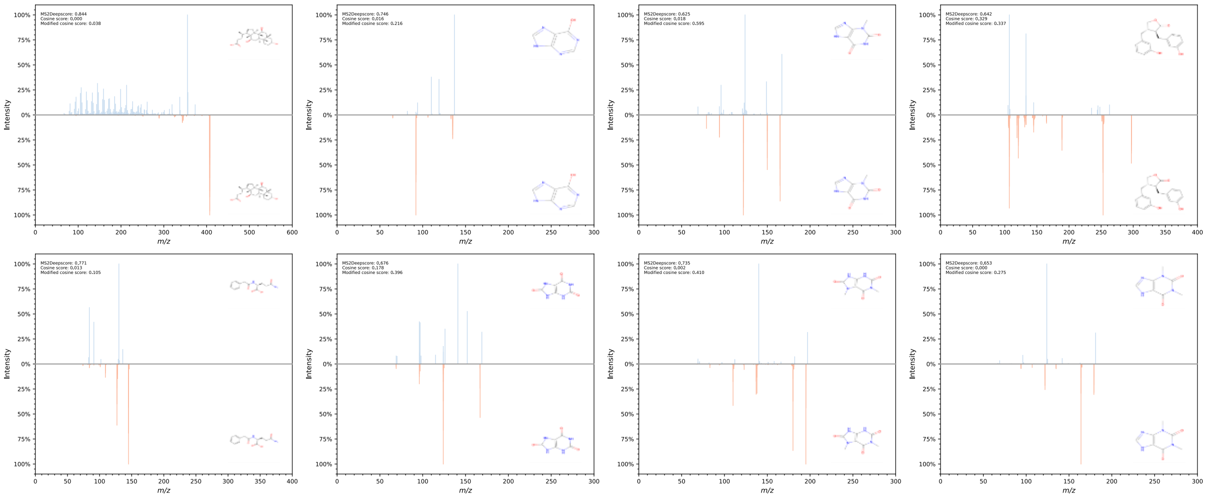

***Supplementary Figure 10.2: Spectrum comparisons of positive and negative ionization mode spectra from the urine case study.*** *Structures are putatively annotated through MS2Query predictions and all have a MS2Query score of at least 0.8 and a mass difference < 0.1 for the precursor m/z. Spectrum pairs are selected for which a positive and negative ionization mode spectrum was detected, with identical annotation. Positive ionization mode spectra are blue and negative ionization mode spectra are orange. The cosine score, the modified cosine score and the MS2DeepScore prediction are given between each spectrum pair.*

#### Supplementary Tables

***Supplementary Table 1: Case study annotations.*** *[*]Levels of confidence in the assignment of the identified metabolites according to the Metabolomics Standards Initiative[2]. Level 1 annotations are confirmed with an in-house standard. Level 2 annotations are confirmed by comparison to a standard in the NIST 23 (2023) mass spectral library[3]. Level 3 annotations are putative annotation based on the MS/MS spectrum interpretation. [**]Spectra with the same cluster number are connected in the molecular network in Figure 5.2.*

| ***m/z*** | **RT in min** | **ionization mode** | **Name** | **Confidence level of annotation*** | **Cluster**** |
|---|---|---|---|---|---|
| 181,0379 | 1,83 | negative | 1-Methyluric acid | 1 | 1 |
| 181,0366 | 1,82 | negative | 1-Methyluric acid | 1 | 1 |
| 181,0385 | 1,83 | negative | 1-Methyluric acid | 1 | 1 |
| 195,0569 | 2,29 | negative | 1,3-Dimethyluric acid | 2 | 1 |
| 195,0524 | 2,61 | negative | 1,7-Dimethyluric acid | 1 | 1 |
| 195,0525 | 2,29 | negative | 1,3-Dimethyluric acid | 2 | 1 |
| 181,0727 | 2,86 | positive | Theophylline | 1 | 1 |
| 181,0725 | 1,11 | positive | 5-Acetylamino-6-amino-3-methyluracil | 2 | 1 |
| 195,091 | 3,61 | positive | Caffeine | 1 | 1 |
| 195,0993 | 3,6 | positive | Caffeine | 1 | 1 |
| 181,074 | 1,11 | positive | 5-Acetylamino-6-amino-3-methyluracil | 2 | 1 |
| 181,0727 | 2,86 | positive | Theophylline | 1 | 1 |
| 195,091 | 3,61 | positive | Caffeine | 1 | 1 |
| 195,0993 | 3,6 | positive | Caffeine | 1 | 1 |
| 181,0725 | 1,11 | positive | 5-Acetylamino-6-amino-3-methyluracil | 2 | 1 |
| 181,074 | 1,11 | positive | 5-Acetylamino-6-amino-3-methyluracil | 2 | 1 |
| 158,0820 | 3,01 | negative | N-Acetyl-L-valine | 3 | 2 |
| 160,0395 | 5,07 | negative | Indole-3-carboxylic acid | 2 | 2 |
| 190,0499 | 3,20 | positive | Kynurenic acid | 1 | 2 |
| 162,0557 | 4,11 | positive | 2,8-Quinolinediol | 2 | 2 |
| 162,0564 | 4,11 | positive | 2,8-Quinolinediol | 2 | 2 |
| 162,0538 | 5,15 | positive | Indole-3-carboxylic acid | 2 | 2 |
| 190,0520 | 3,21 | positive | Kynurenic acid | 1 | 2 |
| 190,0499 | 3,20 | positive | Kynurenic acid | 1 | 2 |
| 162,0557 | 4,11 | positive | 2,8-Quinolinediol | 2 | 2 |
| 162,0564 | 4,11 | positive | 2,8-Quinolinediol | 2 | 2 |
| 162,0538 | 5,15 | positive | Indole-3-carboxylic acid | 2 | 2 |
| 190,0468 | 3,11 | positive | Kynurenic acid | 1 | 2 |
| 190,0468 | 3,11 | positive | Kynurenic acid | 1 | 2 |
| 190,0498 | 3,15 | positive | Kynurenic acid | 1 | 2 |
| 190,0498 | 3,15 | positive | Kynurenic acid | 1 | 2 |
| 190,0520 | 3,21 | positive | Kynurenic acid | 1 | 2 |
| 151,0349 | 2,95 | positive | Homogentisic acid | 2 | 3 |
| 151,0349 | 2,95 | positive | Homogentisic acid | 2 | 3 |
| 151,0351 | 1,33 | negative | 3,4-Dihydroxyphenylglycol | 2 | 3 |
| 407,2797 | 9,7245 | negative | Cholic acid | 1 | UMAP |
| 355,2637 | 9,73 | positive | Cholic acid | 1 | UMAP |

*Supplementary Table 2: Jupyter notebooks used for generating the figure.*

| Figure | Hyperlink to Jupyter notebook |
|---|---|
| 1 | Not generated in a notebook. |
| 2 | https://github.com/matchms/MS2DeepScore/blob/2.5.3/notebooks/model_benchmarking/create_benchmarking_plots.ipynb |
| 3 | Notebooks in https://github.com/matchms/ms2deepscore/tree/2.5.3/notebooks/case_studies_ms2deepscore_2 and visualization in Cytoscape[4]. |
| 4 | https://github.com/matchms/MS2DeepScore/blob/2.5.3/notebooks/model_benchmarking/EmbeddingEvaluator_benchmarking.ipynb |
| SI 1.1 | https://github.com/matchms/MS2DeepScore/blob/2.5.3/notebooks/model_benchmarking/compare_model_trained_on_spectra_with_min_nr_of_peaks.ipynb |
| SI 1.2 - SI 1.7 | https://github.com/matchms/MS2DeepScore/blob/2.5.3/notebooks/model_benchmarking/hyperparameter_optimization.ipynb |
| SI 2 | https://github.com/matchms/MS2DeepScore/blob/2.5.3/notebooks/model_benchmarking/optimize_sampling_algorithm_settings.ipynb |
| SI 3 | https://github.com/matchms/ms2deepscore/blob/2.5.3/notebooks/model_benchmarking/compare_to_ms2deepscore_1/Create_plots.ipynb |
| SI 4 | https://github.com/matchms/MS2DeepScore/blob/2.5.3/notebooks/model_benchmarking/Comparison_to_single_ion_mode_models.ipynb |
| SI 5 | https://github.com/matchms/MS2DeepScore/blob/2.5.3/notebooks/model_benchmarking/Modified_cosine_heatmap.ipynb |
| SI 6 and 7 | https://github.com/matchms/MS2DeepScore/blob/2.5.3/notebooks/model_benchmarking/EmbeddingEvaluator_benchmarking.ipynb |
| SI 9 | ms2deepscore/notebooks/case_studies_ms2deepscore_2/Blood_case_study at 2.5.3 · matchms/ms2deepscore and visualized in Cytoscape[4]. |
| SI 10 | https://github.com/matchms/ms2deepscore/blob/2.5.3/notebooks/case_studies_ms2deepscore_2/Urine_case_study/visualize_example_spectra.ipynb |

*Supplementary Table 3: Availability of raw data for reproducing results in used notebooks.*

| **Notebook** | **Raw data available?** | **Raw input data automatically downloaded?** |
|---|---|---|
| model_benchmarking/ create_benchmarking_plots.ipynb | Yes | Yes |
| model_benchmarking/ EmbeddingEvaluator_benchmarking.ipynb | Yes | Yes |
| model_benchmarking/ Modified_cosine_heatmap.ipynb | Yes | Yes |
| model_benchmarking/ optimize_sampling_algorithm_settings.ipynb | Yes | Yes |
| model_benchmarking /hyperparameter_optimization.ipynb | Yes | Yes |
| case_studies_ms2deepscore_2/Urine_case_study | Yes | Yes |
| model_benchmarking/ compare_model_trained_on_spectra_with_min_nr_of_ peaks.ipynb | No | No |
| model_benchmarking/ comparison_to_single_ion_mode_models.ipynb | No | No |
| https://github.com/matchms/MS2DeepScore/blob/2.5.3/ notebooks/model_benchmarking/compare_to_ms2deeps core_1/ | Yes | Yes |

### References


1. Djoumbou Feunang, Y. et al. ClassyFire: automated chemical classification with a comprehensive, computable taxonomy. *Journal of Cheminformatics* **8**, 61 (2016).
2. Sumner, L.W. et al. Proposed minimum reporting standards for chemical analysis Chemical Analysis Working Group (CAWG) Metabolomics Standards Initiative (MSI). *Metabolomics* **3**, 211-221 (2007).
3. NIST (https://www.nist.gov/programs-projects/nist23-updates-nist-tandem-and-electron-ionization-spectral-libraries; 2023).
4. Shannon, P. et al. Cytoscape: a software environment for integrated models of biomolecular interaction networks. *Genome research* **13**, 2498-2504 (2003).
